# Serum levels of adipokines and insulin are associated with markers of brain atrophy and cognitive decline in the spectrum of Alzheimer’s Disease

**DOI:** 10.1101/2023.09.06.556528

**Authors:** Isabel Garcia-Garcia, Farooq Kamal, Olga Donica, Mahsa Dadar, Alzheimer’s Disease Neuroimaging Initiative

## Abstract

The discovery that metabolic alterations often coexist with neurodegenerative conditions has sparked interest in the examination of gastrointestinal factors as potential modulators of brain health. Here, we examined the role of adipokines (leptin, adiponectin, resistin, and IL6) and insulin on different markers of brain atrophy in participants on the spectrum of Alzheimer’s Disease. We included 566 participants from the Alzheimer’s Disease Neuroimaging Initiative (ADNI) dataset with 1063 follow-up time points (average follow-up: one year); and examined the association between gastrointestinal factors and volumetric MRI values, white matter hyperintensities, and measures of cognitive impairment. Higher leptin, resistin, IL6, and insulin were associated with markers of cerebral atrophy, such as lower total brain volume, or higher ventricular volume. Higher leptin and resistin were also associated with greater impairment in daily life activities. Higher adiponectin was associated with lower ventricle volume. There was no association between adipokines or insulin with white matter hyperintensities. Our findings indicate a co-occurrence between alterations in gastrointestinal factors and in brain volume along the preclinical to clinical spectrum of Alzheimer’s Disease. These results suggest that strategies aimed at promoting metabolic health may positively impact brain health.

## Introduction

Alzheimer’s Disease (AD) is a neurodegenerative disorder whose hallmark pathology is the accumulation of amyloid-β plaques. AD is viewed as a continuum, progressing from sub-clinical subjective cognitive (memory) complaints, to mild cognitive impairment (MCI) and to a final stage of dementia (Jack et al. 2018). Along the continuum, a number of metabolic alterations often co-occur, suggesting the existence of strong links between AD and metabolic dysfunctions. For example, the dementia phase is often accompanied by sudden weight loss, dyslipidemia, inflammation, dysregulated insulin, and hyperglycaemia (Johnson, Wilkins, and Morris 2006; Vinuesa et al. 2021). In addition to these metabolic dysfunctions, alterations in the concentrations of adipokines constitute another metabolic mechanism that might modulate the progress of AD and dementia (Kiliaan, Arnoldussen, and Gustafson 2014). Adipokines are molecules secreted by the peripheral white adipose tissue that exert diverse roles in metabolic and immune health, as well as in brain health (Liu et al. 2022). In the context of brain health, the most studied adipokines are leptin, adiponectin, resistin, and interleukin 6 (IL6). The present study will examine them together with insulin, because of the important role of this last pancreatic peptide in metabolism and in neurodegeneration.

Leptin is a hormone that signals appetite satiation and is secreted by the adipocytes in proportion to the fat stores (Izquierdo et al. 2019). In addition to the adipocytes, small amounts of leptin are also secreted by the hypothalamus. In this brain region, leptin inhibits the secretion of the appetite stimulator neuropeptide Y, and increases the secretion of pro-opio-melanocortin (POM), a satiating factor (Coll, Farooqi, and O’Rahilly 2007). Leptin has beneficial effects in hippocampal neuroplasticity and hippocampal-dependent memory (Forny-Germano, De Felice, and Vieira 2018; Irving and Harvey 2014). Patients with AD consistently show lower levels of blood leptin than cognitively unimpaired controls (García-García et al. 2023). Moreover, studies on the Rotterdam and on the Framigham cohorts have shown that higher levels of leptin were protective against dementia conversion after more than 8 years of follow-up (Mooldijk, Ikram, and Ikram 2021; Lieb et al. 2009).

In the context of abdominal obesity, aging, and neurodegeneration, there is often a decrease in the capacity of the body to respond to leptin, a process called leptin resistance. Leptin resistance has been hypothesized to impair protein clearance in AD (Forny-Germano, De Felice, and Vieira 2018). Moreover, leptin resistance might also facilitate chronic inflammation (Martin, Qasim, and Reilly 2008). When levels of leptin are consistently high, leptin creates a pro-inflammatory effect by increasing the proliferative activity of macrophages and by stimulating the secretion of the cytokines interleukin 6 (IL6), interleukin 1 beta (IL1β), and tumor necrosis factor alpha (TNFα) (Correale and Marrodan 2022; Martin, Qasim, and Reilly 2008). Due to these proinflammatory effects, high concentrations of leptin are largely regarded as detrimental for cerebrovascular disease, and can contribute to the formation of the atherosclerotic plaques (Liu et al. 2022; Reilly et al. 2004).

Adiponectin is an anti-inflammatory peptide secreted exclusively in the white adipose tissue. It inhibits the maturation and proliferation of macrophages and the secretion of TNFα and interferons, while at the same time it increases the secretion of anti-inflammatory cytokines, such as TNFβ and IL10 (Correale and Marrodan 2022). Low adiponectin has been associated with increases in cardiometabolic burden, with type 2 diabetes mellitus, and with abdominal obesity (Forny-Germano, De Felice, and Vieira 2018). Adiponectin protects against the formation of the atherosclerotic plaques (Liu et al. 2022). However, the association between adiponectin and AD is still unclear. Animal studies suggest that adiponectin protects against AD pathology (Forny-Germano, De Felice, and Vieira 2018), while human longitudinal studies have not found so far consistent associations between low adiponectin and risk of AD (García-García et al. 2023).

Resistin is secreted by macrophages, which in the context of abdominal obesity are infiltrated in large numbers in the adipose tissue. Resistin stimulates the secretion of the pro-inflammatory cytokines TNFα, IL6, and IL1β, and promotes insulin resistance and systemic chronic inflammation (Correale and Marrodan 2022). Resistin also exacerbates the development and worsening of atherosclerosis (Liu et al. 2022).

IL6 is a proinflammatory cytokine that is secreted by several cell types, including adipocytes. Specifically, the adipocyte secretion of IL6 tends to promote systemic inflammation, as seen in some cardiometabolic diseases such as T2DM and obesity (Luan et al. 2023). Furthermore, IL6 is a contributor to the atherosclerotic plaques (Liu et al. 2022).

Insulin is a pancreatic hormone secreted by the β-cells that regulates the cellular uptake of glucose. When the blood levels of insulin are high, cells enhance their glucose uptake, lowering the levels of glucose in the bloodstream. When the blood concentrations of insulin are chronically high, the cells develop insulin resistance, meaning that their capacity to respond to insulin becomes compromised. Similar to adipokines, insulin resistance might affect brain health by two main mechanisms. First, insulin affects protein deposition. In a homeostatic state, insulin protects against amyloid-β toxicity and by promoting protein clearance (Kellar and Craft 2020). By contrast, insulin resistance in the context of AD has been associated with higher deposition of amyloid-β protein in fronto-temporal regions (Willette et al. 2015) and with synaptic impairment (Forner et al. 2017). Second, insulin resistance, and the subsequent chronic hyperinsulinemia, stimulates vasoconstriction (Kellar and Craft 2020), while insulin resistance has been associated with cerebral hypoperfusion (Cui et al. 2017). In agreement with the negative effect of insulin resistance on the brain, type 2 diabetes mellitus is a risk factor for AD and other dementias (Yu et al. 2020; Livingston et al. 2020).

Previous studies have examined the association between blood levels of adipokines and the incidence of mild cognitive impairment or AD (García-García et al. 2023; Kim et al. 2022; Mooldijk, Ikram, and Ikram 2021). However, these same studies are generally limited in their examination of neurological and cognitive correlates of neurodegeneration (García-García et al. 2023). In this paper, we provide a comprehensive overview of different proinflammatory and antiinflammatory adipokines and insulin in relation with several markers of brain senescence and cognitive impairment in participants in the spectrum of AD.

## Methods

### Alzheimer’s Disease Neuroimaging Initiative

The data used in this article was obtained from the ADNI database (adni.loni.usc.edu), which was established as a joint public-private initiative led by Michael W. Weiner, MD, in 2003. The database’s primary objective is to investigate whether a combination of magnetic resonance imaging (MRI), positron emission tomography (PET), other biological markers, and clinical and neuropsychological assessments can be used to measure the progression of MCI and AD. The study received ethical approval from the review boards of all the institutions involved, and written consent was acquired from the participants or their study partner. The study’s participants were selected from all the ADNI cohorts (ADNI-1, ADNI-2, ADNI-GO, and ADNI-3).

### Participants

Participant inclusion and exclusion criteria for this study can be found at www.adni-info.org. To be included in the study, participants had to be between the ages of 55 and 95 and not exhibit any symptoms of depression. Healthy older adults had no evidence of memory decline or impaired global cognition, as assessed by the Wechsler Memory Scale, Mini-Mental Status Examination (MMSE), and Clinical Dementia Rating (CDR). Participants with MCI had scores between 24 and 30 on the MMSE, a CDR score of 0.5, and abnormal scores on the Wechsler Memory Scale. Individuals with AD had abnormal memory function on the Wechsler Memory Scale, MMSE scores between 20 and 26, a CDR of 0.5 or 1.0, and met the criteria of the National Institute of Neurological and Communicative Disorders and Stroke and the Alzheimer’s Disease and Related Disorders Association (Tierney et al. 1988).

### Plasma Data

The present study analyzed plasma adiponectin, leptin, resistin, insulin, and IL6 data from the Biomarkers Consortium Project from ADNI. ADNI Researchers used a multiplex immunoassay panel based on the Luminex xMAP platform from Rules-Based Medicine (RBM) in Austin, TX, to analyze the adipokines in plasma. The platform from RBM is an immunological method that can detect and quantify multiple target proteins simultaneously by detecting fluorescent microspheres. Methods for quantifying the data are available at http://adni.loni.usc.edu/wp-content/uploads/2010/11/BC_Plasma_Proteomics_Data_Primer.pdf.

A total of 566 participants with Adipokines measures available were included in this study [58 healthy controls (NC), 396 MCI, and 112 AD]. The groups had the following number of average follow-up times: 1) NC: 0.95 years, 2) MCI: 0.88 years, 3) AD: 0.87 years.

### Structural MRI acquisition and processing

MRI scanning for all participants in the study followed standardized acquisition protocols developed and implemented by ADNI. Additional information about the MRI protocols and imaging parameters can be found at http://adni.loni.usc.edu/methods/mri-tool/mri-analysis/. MRI data for all participants were obtained from the ADNI public website.

Using our standard pipeline, pre-processing of all T1w scans included noise reduction (Coupe et al. 2008), intensity inhomogeneity correction (Sled, Zijdenbos, and Evans 1998), and intensity normalization into range [0-100]. The pre-processed images were then linearly (9 parameters: 3 translation, 3 rotation, and 3 scaling) (Dadar, Fonov, et al. 2018) registered to the MNI-ICBM152-2009c average template (Fonov et al. 2011).

### WMH measurements

Our previously validated and well-established methods were employed to obtain measurements of white matter hyperintensities (WMHs) (Dadar et al. 2017). This method has been used in other aging and neurodegenerative disease multi-center cohort studies (Anor et al. 2021; Dadar et al. 2020), including the ADNI (Kamal et al. 2023b, [a] 2023). To segment WMHs, the method utilizes a combination of the T1w contrast and features related to the location and intensity of manually segmented scans from a representative library (50 ADNI participants that were not part of the study) (Dadar et al. 2017). The segmentations were automatically performed in combination with a random forest classifier to detect the WMHs in new images. The study used T1w images to segment WMHs instead of FLAIR and T2w/PD scans, as ADNI1 only acquired T2w/PD images at a resolution of 1×1×3 mm^3^ whereas ADNI2/GO acquired only axial FLAIR images with resolutions of 1×1×5 mm^3^, and ADNI3 acquired sagittal FLAIR images with resolutions of 1×1×1.2 mm^3^. There are potential inconsistencies in WMH measurements across the ADNI cohorts. Thus, we opted to include consistently acquired T1w images to measure WMH burden. T1w-based WMH volumes have been previously shown to be highly correlated with FLAIR and T2w based WMH loads in the ADNI dataset (Dadar, Maranzano, et al. 2018; Dadar et al. 2020). An experienced rater assessed the quality of registrations and WMH segmentations and cases that did not pass the quality control were excluded from analyses (N = 59). WMH load was determined as the standard space volume of all voxels identified as WMH and was thus normalized for head size. WMH volumes were calculated based on Hammers Atlas (Dadar et al. 2017; Dadar, Maranzano, et al. 2018). Normalization was achieved through log-transformation of all WMH volumes.

A previously validated patch-based label fusion segmentation technique was used to segment the ventricles based on T1w images (Coupé et al. 2011; Manera et al. 2022). Hippocampal, entorhinal cortex, and medial temporal lobe volumes were assessed using the volumes provided by ADNI.

### Cognitive Scores

Participant cognitive scores included a measure of difficulties in activities of daily living using the Functional Activities Questionnaire (FAQ), a measure of global cognitive functioning using the Clinical Dementia Rating – Sum of Boxes (CDR-SB), a measure of episodic memory using Rey Auditory Verbal Learning test (RAVLT), and a measure of executive functioning using the Digit Span Test Score (DIGITSCOR). These specific tests were chosen because more than 80% of the participants completed them. These cognitive scores were downloaded from the ADNI website.

### Statistical Analysis

T-tests and chi-square analyses were performed on the demographic information and baseline measurements, corrected for multiple comparisons using Bonferroni correction. A series of longitudinal mixed effects models were used to assess the relationship between adipokines and insulin and risk factors including a history of hypertension, the modified Hachinski ischemic score, diabetes, and BMI, with the adipokine measures as dependent variables, and hypertension, Hachinski score, diabetes, and BMI as independent fixed variables of interest. The models included age, years of education, sex, and baseline diagnosis as covariates. Similarly, longitudinal mixed effects models were used to assess the relationship between adipokines, brain, and cognitive measurements. Adipokines were considered as continuous independent fixed variables of interest and cognitive (e.g. CDRSB score) and brain (e.g. whole brain volume) measurements were considered as continuous dependent variables. All models included age, sex, years of education, baseline diagnosis, and BMI as covariates. Participant IDs were included as categorical random effects. All continuous values were z-scored within the population prior to the regression analyses. All results were corrected for multiple comparisons using false discovery rate (FDR) controlling technique with a significance threshold of 0.05, p-values are reported as raw values with significance then determined by FDR correction (Benjamini and Hochberg 1995). To complete the longitudinal assessment, we used function fitlme in MATLAB. All statistical analyses were performed using MATLAB version 2021a.

## Results

Table 1 summarizes the demographic and clinical information of the participants included in this study. Differences in demographics and baseline characteristics are reported only for those that survived correction for multiple comparisons. As expected, the AD group had significantly higher CDRSB scores and more APOE4+ individuals than MCI and controls. The MCI group had significantly higher CDRSB scores and more APOE4+ individuals than the controls (p < 0.001). The AD group also had significantly higher resistin levels than the MCI group (p = 0.0001). No significant differences were observed in the other demographics and baseline characteristics variables. Figure 1 shows the correlations between the different adipokine levels and BMI.

**Table 1.** Demographic information for the participants included in this study. MCI = Mild Cognitive Impairment; AD = Alzheimer’s Dementia; BMI = Body Mass Index; CDRSB = Clinical
Dementia Rating Sum of Boxes. Note: Statistically significant results are reported if they survive correction
for multiple comparisons using the Bonferroni method. ^ Significantly different from control. # significantly
different from MCI.

| Measure | Cognitively unimpaired participants | Participants with MCI | Participants with AD |
| --- | --- | --- | --- |
| N (N Female) | 58 (28) | 396 (140) | 112 (47) |
| Baseline Age | 75.06 ± 5.76 | 74.70 ± 7.39 | 74.75 ± 80.8 |
| Education | 15.67 ± 2.78 | 15.64 ± 3.04 | 15.03 ± 3.19 |
| BMI | 27.24 ± 4.11 | 26.07 ± 3.94 | 25.53 ± 3.02 |
| N Diabetes (%) | 2 (3.45%) | 27 (6.82%) | 5 (4.46%) |
| Hypertension (No / Yes) | 29 / 29 | 202 / 194 | 53 / 59 |
| APOE4 (0/1/2) | 53 / 5 / 0 | 185 / 164 / 47 | 36 / 53 / 23 |
| Baseline CDRSB | 0.02 ± 0.11 # | 1.60 ± 0.88 ^ | 4.32 ± 1.56 #^ |
| Baseline adiponectin | 0.70 ± 0.29 | 0.76 ± 0.25 | 0.80 ± 0.25 |
| Baseline insulin | 0.37 ± 0.32 | 0.32 ± 0.31 | 0.30 ± 0.29 |
| Baseline leptin | 1.10 ± 0.42 | 0.92 ± 0.40 | 0.96 ± 0.45 |
| Baseline resistin | 0.51 ± 0.12 | 0.47 ± 0.16 | 0.54 ± 0.14# |
| Baseline interleukin 6 | 1.50 ± 0.11 | 1.46 ± 0.14 | 1.46 ± 0.11 |

**Figure 1.**
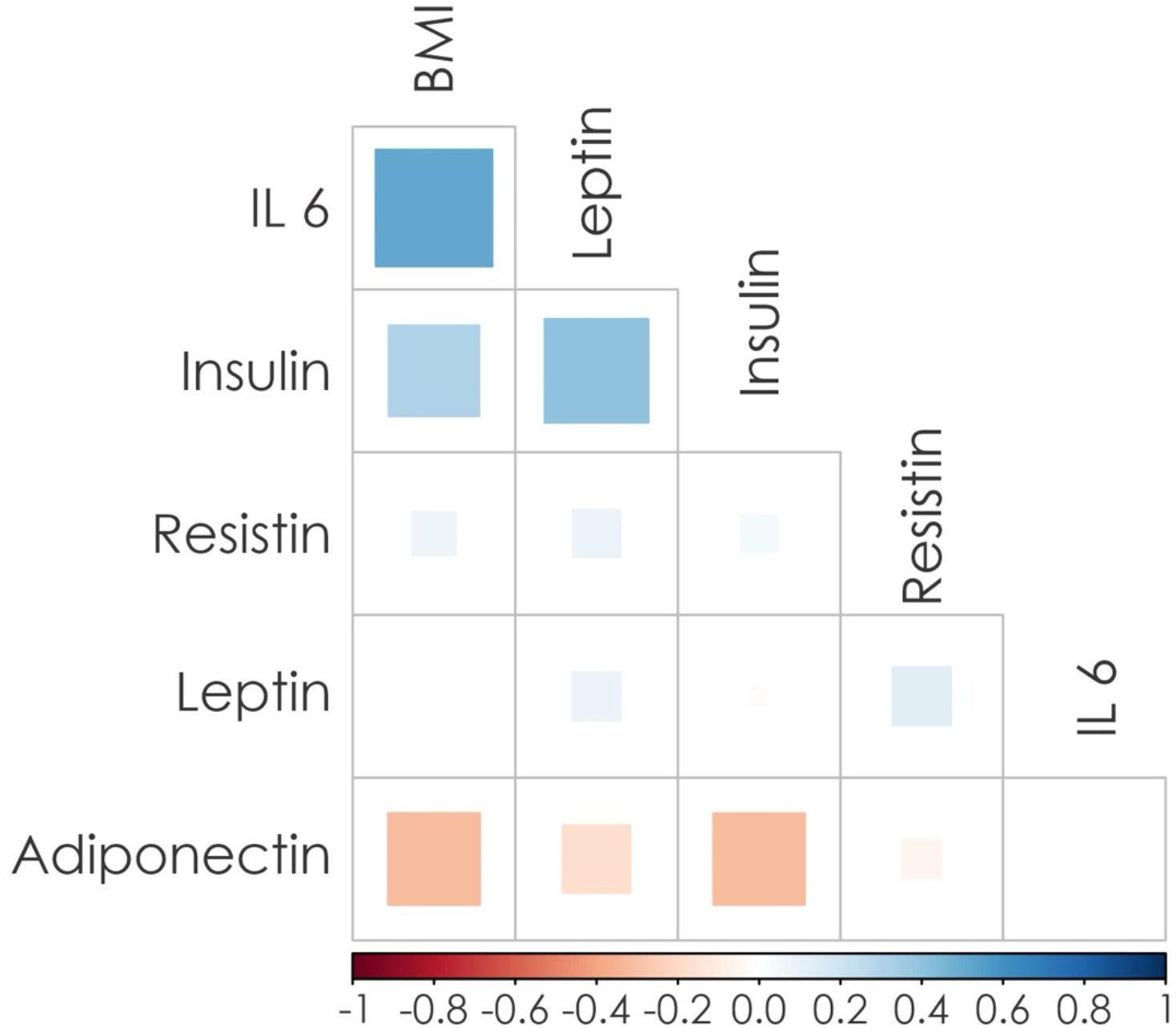
Correlations between the different adipokines, insulin, and body mass index (BMI).

Table 2 summarizes the results of the longitudinal mixed effects models investigating the impact of a history of hypertension, Hachinski ischemic score, diabetes, and obesity on the adipokine and insulin measurements. Hypertension and greater Hachinski scores were both significantly associated with higher insulin (t = 3.67, 3.12 respectively; p<0.001) and leptin (t = 5.76, 3.90 respectively; p<0.001) levels. As expected, diabetes was also significantly associated with higher insulin levels (t = 2.89 ; p=0.003). Finally, higher BMI values were associated with lower adiponectin (t = -8.31 ; p<0.001) and higher insulin (t = 3.03 ; p<0.001) and leptin (t = 23.60 ; p<0.001) levels.

**Table 2.** Summary results of the mixed effects models investigating the impact of risk factors on adipokine measures.

| Measure | Adiponectin | Insulin | Leptin | Resistin | Interleukin 6 |
| --- | --- | --- | --- | --- | --- |
| Hypertension | -0.71 | **3.67\*\*** | **5.76\*\*** | 1.71 | 0.59 |
| Hachinski score | -0.52 | **3.12\*\*** | **3.90\*\*** | 0.66 | 0.51 |
| Diabetes | 0.20 | **2.89\*** | 1.84 | -0.51 | -0.78 |
| BMI | **-8.31\*\*** | **8.03\*\*** | **23.60\*\*** | 1.55 | 0.59 |

\* $p < 0.01$ ; \*\* $p < 0.001$

Table 3 summarizes the results of the longitudinal mixed effects models, assessing the association between adipokines, brain, and cognitive measures. Higher adiponectin levels were significantly associated with lower ventricle volumes (t = -4.534; p<0.001), indicating a positive impact of adiponectin on overall brain health. Similarly, higher insulin and IL6 levels were also associated with greater ventricle volumes (t = 3.021; p<0.01 for insulin and t = 3.158; p<0.001 for IL6) and lower total brain volumes (t = -2.567; p<0.01 for insulin and t = -3.079; p<0.01 for IL6), indicating a negative impact of higher insulin and IL6 levels on brain health. Higher leptin levels were associated with higher FAQ scores (t = 2.416; p<0.01), higher CDRSB scores (t = 3.198; p<0.001), greater ventricle volumes (t = 2.495; p<0.01), and lower total brain (t = -3.546; p<0.001) and hippocampal volumes (t = -2.737; p<0.01), indicating a negative impact of higher leptin levels on overall brain health, hippocampal atrophy, global cognition, and functional activities of daily living. Finally, higher resistin levels were associated with higher FAQ scores (t = 3.227; p<0.001), and lower hippocampal (t = -3.066; p<0.01) and entorhinal cortex (t = -2.848; p<0.01) volumes, indicating a negative impact of resistin on more AD-related neurodegeneration and functional activities of daily living (Figure 2). The models were also repeated with hypertension, Hachinski score, and diabetes as additional covariates, yielding similar results in terms of effect size and significance.

**Table 3.** Summary results of the mixed effects models investigating the impact of adipokines on brain and cognitive measures.

| Measure | Adiponectin | Insulin | Leptin | Resistin | Interleukin 6 |
| --- | --- | --- | --- | --- | --- |
| Functional activities (FAQ) | -0.407 | 1.559 | **2.416\*** | **3.227\*\*** | -0.033 |
| Global cognition (ADAS13) | -0.999 | 1.708 | 1.871 | 0.486 | -0.372 |
| Global cognition (CDRSB) | -1.143 | 2.142 | **3.198\*\*** | 1.102 | 0.301 |
| Memory: RAVLT_immediate | 0.817 | -1.510 | -1.353 | -2.234 | 0.748 |
| Memory: RAVLT_learning | 0.429 | -0.126 | -0.973 | -1.524 | -0.183 |
| Memory: RAVLT_forgetting | 0.085 | -0.852 | 1.146 | -0.960 | 0.172 |
| Executive function (Digit Score) | -0.858 | 0.385 | -1.097 | -0.829 | -0.109 |
| Hippocampus volume | 2.241 | -0.689 | **-2.737\*** | **-3.066\*** | -0.805 |
| Whole Brain Volume | 1.987 | **-2.567\*** | **-3.546\*\*** | -0.685 | **-3.079\*** |
| Ventricle volume | **-4.534\*\*** | **3.021\*** | **2.495\*** | 1.609 | **3.158\*\*** |
| WMH volume | -0.806 | 1.566 | 1.187 | -1.005 | -0.346 |
| Entorhinal cortex volume | 1.079 | -0.463 | -0.209 | **-2.848\*** | 0.566 |
| Medial Temporal lobe volume | 0.905 | -1.449 | -1.742 | -1.900 | -1.452 |
| CSF Amyloid beta | 0.049 | 0.281 | 0.867 | -0.789 | -0.653 |
| CSF Tau | 0.734 | 1.381 | -0.629 | 0.990 | 0.126 |

\* $p < 0.01$ ; \*\* $p < 0.001$

**Figure 2.**
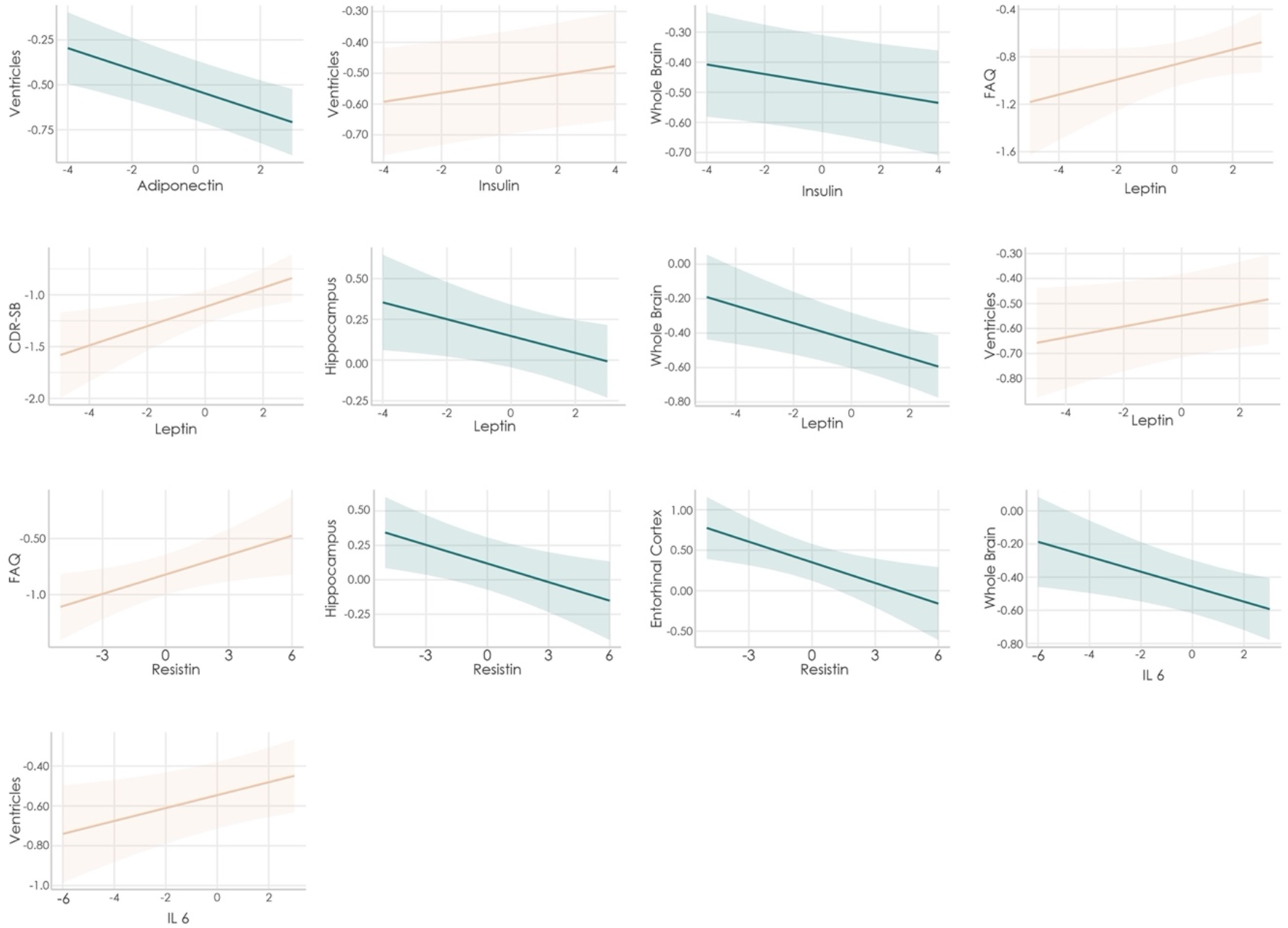
Significant associations between adipokine values and brain outcomes in participants in the spectrum of Alzheimer’s disease.

## Discussion

This study examines the neurobehavioral correlates of serum levels of adipokines and insulin in participants with diverse gradients of AD pathology and across longitudinal follow-ups. Our results show that higher leptin, IL6, and insulin were associated with global markers of brain atrophy, such as whole brain volume or ventricular volume. Participants with higher levels of leptin and resistin had lower gray matter volume in temporal-medial structures along with greater impairment in functional activities. Finally, higher levels of adiponectin were correlated with less ventricular volume. Our findings align well with the hypothesis that metabolic factors, such as adipokines and insulin, are implicated in brain senescence and neurodegeneration. They further hint at the possibility that strategies to boost metabolic health may also have important positive effects on brain health. At the same time, the non-interventional nature of the study precludes us from uncovering causal mechanisms between these gastrointestinal factors and brain integrity.

Among the gastrointestinal factors examined, we found that leptin was associated with the greatest number of unfavorable brain outcomes, including reduced global brain volume, lower hippocampal volume, and lower cognitive scores. These results are consistent with previous studies linking elevated leptin levels to increased risk of cognitive decline and neurodegeneration (Forny-Germano, De Felice, and Vieira 2018). Similar to leptin, higher levels of resistin were negatively associated with impairment in daily function and with reduced hippocampal and entorhinal cortex volumes, suggesting a link between resistin and AD-related neurodegeneration. These associations remained after accounting for hypertension, Hachinski score, and diabetes as additional covariates, reinforcing the independent contributions of adipokines to brain health and cognitive outcomes.

Adipokines have important effects on systemic inflammation. High levels of the proinflammatory adipokines leptin, resistin, and IL6, facilitate a state of systemic chronic inflammation (Liu et al. 2022), and this state is also promoted by insulin resistance. Chronic inflammation jeopardizes the integrity of the blood brain barrier (BBB) and enhances its permeability, leading to increases in the concentrations of pro-inflammatory signals in the brain tissue (Walker et al. 2022). Chronic inflammation can impair microglial function, impair its ability to clear protein deposition, and contribute this way to brain atrophy and neurodegeneration (Walker et al. 2022). Similarly, we found that higher blood levels of pro-inflammatory adipokines and higher concentrations of insulin were associated with lower brain tissue, as indicated by lower cerebral brain volume and higher ventricular space volume. Conversely, adiponectin, an anti-inflammatory adipokine, was associated with lower ventricle volume, which suggests the possibility that adiponectin has protective effects in the brain.

We have focused on the association between gastrointestinal factors and markers of cerebral atrophy in participants with AD pathology. The idea that metabolic and brain factors go hand-by-hand aligns with the possibility that strategies to improve metabolic health might also benefit brain health. It has been shown that moderate-to-vigorous physical activity has beneficial effects on cognition and that it is associated with reduced risk of cognitive impairment (Erickson et al. 2019). Moreover, results from the FINGER trial, a lifestyle intervention that included diet counseling, physical training, cognitive training, and monitoring of vascular risk factors, showed that this multifactorial intervention had beneficial cognitive effects, especially regarding executive functions and cognitive speed (Ngandu et al. 2015).

This paper has some methodological limitations. First, individuals in the ADNI dataset may not be representative of other parts of the population, and only a subset of participants had measurements of adipokines or insulin available. Second, adipokines and insulin were measured in the blood, which provides at best an indirect estimation of the CNS levels of these gastrointestinal factors. Third, the lack of extended follow-up periods (>5 years), does not allow us to detect the long-term longitudinal impacts of gastrointestinal factors on brain and cognition trajectories.

In conclusion, our findings highlight the interplay between gastrointestinal factors, specifically adipokines and insulin, and neurocognitive in the context of Alzheimer’s Disease pathology. Higher leptin, resistin, IL6, and insulin were associated with global brain atrophy markers, indicative of potential neurodegeneration. Conversely, elevated adiponectin was linked to reduced signs of ventricular enlargement, suggestive of a potential positive association with brain integrity. Our findings support the possibility that bolstering metabolic health might additionally benefit brain health in participants with underlying pathology of AD.

## Notes

### Competing Interest Statement

IGG and OD are currently employed at Clinique la Prairie

